# Clonal morphology-guided combination therapies overcome heterogeneity-driven drug tolerance

**DOI:** 10.64898/2026.01.31.703021

**Authors:** Xiaoxi Li, Ling Liu, Yuanyuan Yan, Mengfang Yang, Lingli Luo, Yong Jiang, Weijun Ren

**Author notes:** These authors contributed equally: Xiaoxi Li, Ling Liu. Correspondence (Xiaoxi Li).

## Abstract

Tumor heterogeneity plays a critical role in tumor relapse and the development of drug resistance. Current personalized strategies, based on genetic profiling or bulk tumor drug sensitivity testing, offer limited clinical value as they overlook clonal heterogeneity. Here, through establishing primary tumor cell cultures from multiple metastatic sites, we observed that tumor cells exhibit high morphological plasticity and readily form distinct clonal morphologies during monoclonal expansion. Moreover, these heterogeneous clones displayed distinct tumorigenic and histological traits in orthotopic models, which may imply their differing clinical relevance in tumor progression. We also found that clonal morphological heterogeneity was exhibited by both established cell lines and primary lines derived from chemically induced tumor models. Subsequently, we analyzed the drug sensitivity profiles of tumor clones and identified clone-specific sensitive drugs. Despite limited *in vitro* synergy in pairwise combinations, both *in vitro* and *in vivo* assays confirmed their potent, clone-selective inhibition, suggesting rational drug combinations can precisely target distinct clonal populations. Taken together, Collectively, our findings propose that a heterogeneity-informed drug selection paradigm, enabled by rapid clonal morphology analysis of patient-derived cells, can be used to prevent recurrence and overcome resistance.

**Graphic Abstract:** 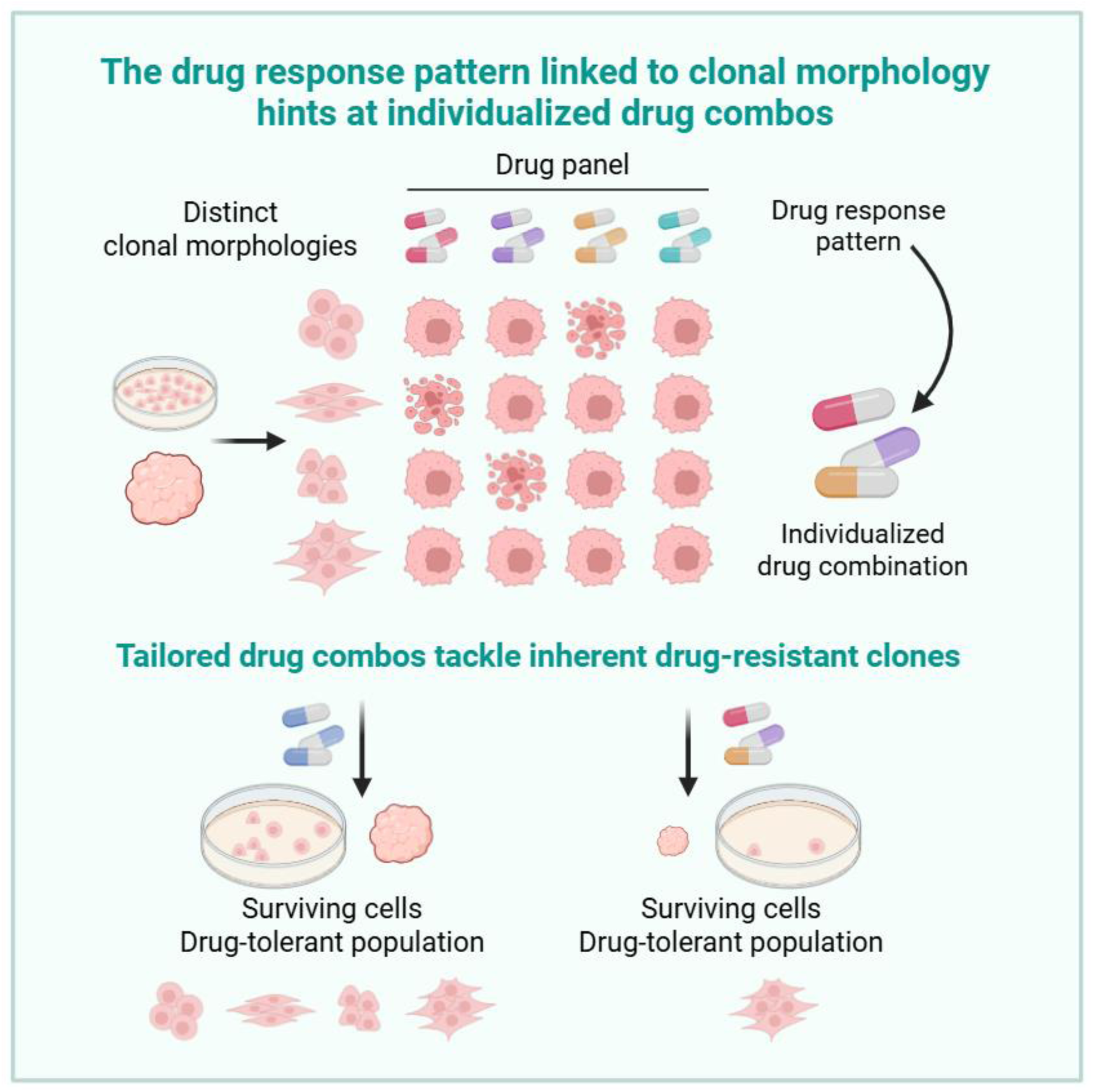

**Highlights:**

1. Tumor cells develop distinct morphologies during clonal expansion.
2. Distinct morphologies serve as simple markers of tumor heterogeneity.
3. Distinct clonal morphology defines a unique drug-response profile.
4. Clone-selective drug combinations combat heterogeneity-driven resistance.

## Introduction

With advances in cancer genomics, the genetic profiles of patient tumors have become a critical foundation for personalized cancer therapy, offering essential guidance for molecular stratification and treatment selection. However, relying solely on genetic variation information is insufficient to achieve the goals of personalized precision treatment, dynamic monitoring, and drug efficacy evaluation. Tumors with identical genetic profiles can display vastly different clinical behaviors in growth, metastasis, and drug response, dictated by distinct epigenetic landscapes^1^. The concept of Functional Precision Medicine (FPM) has gained widespread acceptance in recent years^2–5^. This approach directly utilizes viable tumor tissue samples from patients for drug sensitivity testing—rather than relying solely on genetic data—to predict anti-tumor drug efficacy and guide personalized treatment. Several clinical studies have demonstrated that treatment plans based on FPM results can improve patient outcomes^6–8^. Nevertheless, tumor tissue is composed of highly heterogeneous clones that evolve dynamically. This intra-tumoral heterogeneity is a primary driver of tumor recurrence and intrinsic drug resistance. Current FPM strategies, while personalized, are predominantly based on inter-patient genetic variation. Consequently, the derived drug sensitivity profiles provide only a generalized tumor response map. They overlook the critical differences in drug sensitivity among distinct intra-tumoral clones and the therapeutic resistance masked by this inter-clonal heterogeneity.

Combination therapy is a major advance in oncology, aiming to lower doses, reduce toxicity, and forestall resistance compared to single agents^9^. However, the current paradigm for discovering combinations focuses predominantly on identifying drug synergy^10^, failing to consider the critical influence of tumor heterogeneity. Although synergy can improve efficacy and safety, it cannot overcome resistance rooted in heterogeneity. Such resistance is either acquired or intrinsic. Intrinsic resistance stems from two distinct scenarios^11–15^: (1) tumor-wide tolerance driven by a patient’s unique genetic background (inter-tumoral/inter-patient heterogeneity), or (2) the coexistence of sensitive and resistant cell clones within the same tumor or across different tumors in the same patient (intra-tumoral heterogeneity). Many experimental studies have confirmed the prevalence of intra-tumoral heterogeneity and its profound impact on tumor clonal evolution, metastatic potential, and therapeutic response^16–21^. Moreover, an *in vitro* drug resistance study has identified non-genetic factors, including transcriptional profiles but not genetic mutations, as key drivers of tumor cell heterogeneity^22^.Consequently, the core capability to distinguish heterogeneous clones and define their drug sensitivity profiles is essential for advancing functional precision medicine, enabling personalized treatment, accurate outcome prediction, resistance surveillance, and informed drug combination discovery.

Tumor cell phenotypic heterogeneity, including morphological plasticity, is a key feature of tumors^23–26^. Several studies have documented diverse cellular morphologies within tumor cell lines, which may correlate with various steps in the metastatic cascade^27–31^. However, the translational value of this morphological diversity in clonal populations remains underexplored. Here, we analysis demonstrates that tumor cell clones from distinct metastatic sites exhibit unique and identifiable morphological signatures, most discernible in low-density cultures, which can serve as biomarkers for clonal heterogeneity. Mapping the drug sensitivity profiles of these clones enables the rational design of clone-tailored drug combinations aimed at mitigating resistance and improving therapeutic efficacy. Thus, we propose that this clonal morphology-based framework represents a transformative paradigm for discovering drug combinations and advancing functional precision medicine into clinical practice.

## Results

### Heterogeneous tumor cell clones derived from distinct metastatic sites form unique clonal morphologies

During tumorigenesis, the accumulation of genetic alterations gives rise to a highly heterogeneous cell population within primary tumors. In contrast, cells at metastatic sites often originate from a single clone, representing a monoclonal outgrowth of a malignant, metastatic-competent subpopulation from the heterogeneous primary tumor^32–37^. Consequently, analyzing the characteristics of these metastatic clones is crucial for understanding tumor recurrence and drug resistance, and for developing preventive strategies.

In this study, we employed an orthotopic transplant and resection strategy to model post-surgical recurrence and metastasis^38,39^. This strategy was designed to clinically mimic primary tumor removal and provide a prolonged timeframe for metastatic development (Fig. 1A). Notably, to enrich for clones with higher metastatic potential than the parental cells at secondary sites, we utilized the EO771 cell line, which has a relatively low metastatic potential, instead of the highly metastatic 4T1 line for orthotopic model establishment. The primary tumor was surgically resected when it reached 400-600 mm³ (Fig. 1B). Recipient mice were monitored and euthanized upon the development of moribund symptoms, with a median survival of 52 days (Fig. 1C). Dissection and analysis of metastatic sites revealed that the majority of recipient mice developed lymph node (6/7) and intestinal (4/7) metastases. A smaller subset exhibited lung metastases (3/7), thoracic colonization (1/7), liver metastasis (1/7), and ureter metastasis (1/7) (Fig. 1D-E). Furthermore, we established five primary cell lines from lymph node and lung metastases of recipient mice #2, #3, and #6. These lines were designated according to the recipient mouse number and metastatic site (prefix “EM” denotes EO771 Metastasis). During culture, we observed that these primary cell lines formed distinctive morphological patterns (Fig. 1F). Three lines—EM2LN (with well-defined, epithelial-like morphology and clear cell borders), EM2Lu (smaller, rounded, stem-cell-like), and EM6LN (lacking distinct morphology and borders, mesenchymal-like)—were selected as heterogeneous tumor clonal models for further expansion and analysis. They represent clones from the same metastatic site in different mice and different sites in the same mouse.

**Fig. 1.**
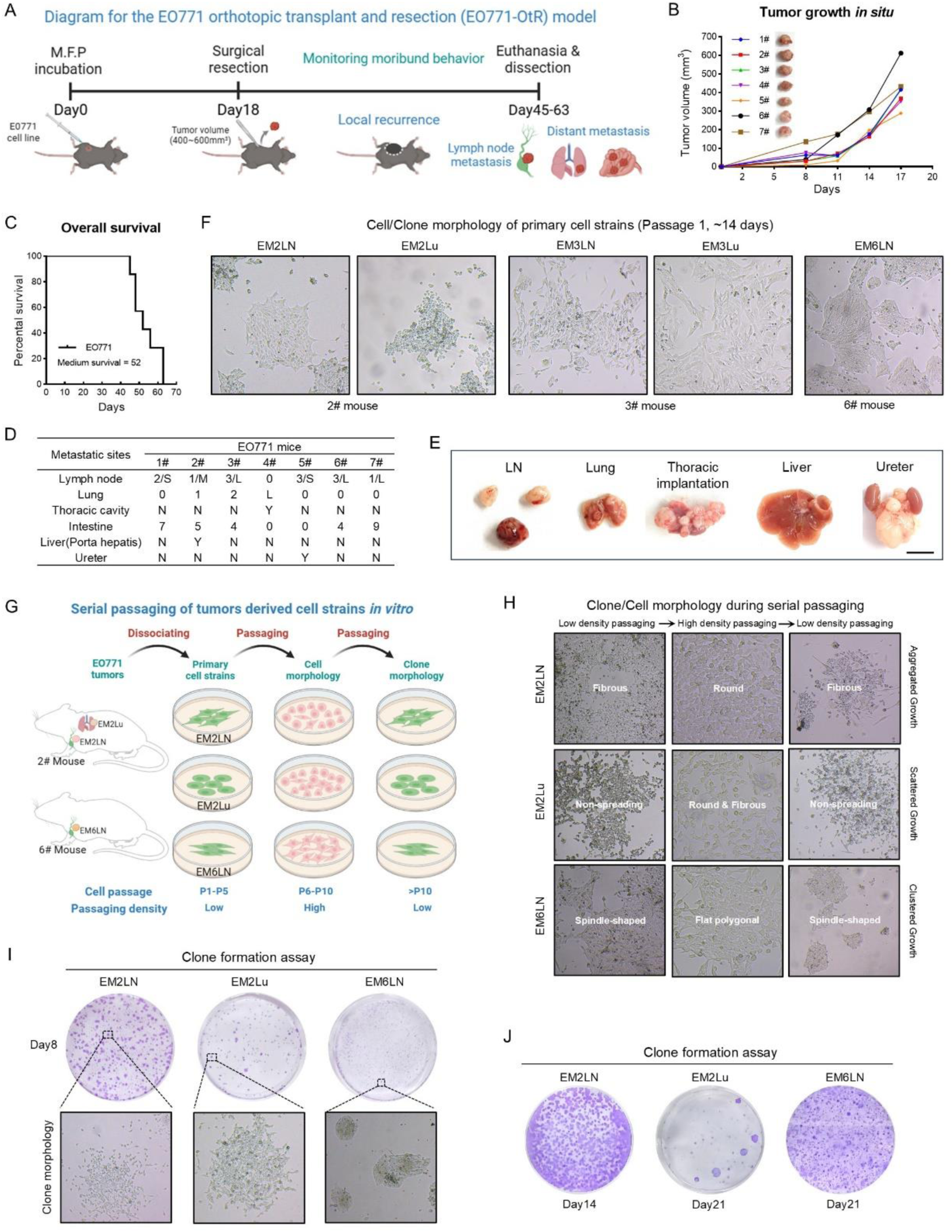
Heterogeneous tumor cell clones possess unique and stable clonal morphologies. A. Schematic of the EO771 orthotopic transplant and resection (OtR) model for studying recurrence and metastasis. B. Growth curve of the primary orthotopic tumors and representative photographs of the surgically resected tumors. C. Kaplan-Meier survival curve of EO771-OtR model mice. D. Statistical analysis of metastatic sites in EO771-OtR mice. E. Representative images of distal metastatic tumors. Scale bar, 8mm. F. Cellular morphology of primary cell lines derived from different metastatic tumors. Scale bar, 200μm. G-H. Morphological features of primary tumor cells and their derived clones during *in vitro* passaging. B Scale bar, 200μm. I. Colony formation and colony morphology of the three EM cell lines in an 8-day colony formation assay. Scale bar, 200μm. J. Colony formation capacity of the three EM cell lines in an extended colony formation assay.

During the culture of heterogeneous tumor cell clones, we observed that their distinct morphological features were stably maintained across passages without loss (Fig. 1G-H). These characteristics showed a density-dependent attenuation, becoming less distinct at high cell density but strikingly prominent in low-density cultures where clonal outgrowths formed (Fig. 1G-H). This persistence indicates that clonal morphology is an intrinsic and heritable property of the tumor cells. Our observations of clonal morphological plasticity are in line with recent reports of highly plastic tumor cells in lung cancer models^40,41^.

To further dissect the behavioral heterogeneity among these clones, we performed colony formation assay. In an 8-day assay, EM2LN formed the largest colonies, EM2Lu produced fewer colonies, while EM6LN generated small colonies that were faintly stained by crystal violet (Fig. 1I). In extended assays, EM2LN exhibited the most rapid growth, forming confluent sheets by day 14. In contrast, EM2Lu and EM6LN grew slower; by day 21, EM2Lu had formed only sparse colonies, and EM6LN colonies remained small (Fig. 1J).

Taken together, we have established heterogeneous tumor cell clones from distinct metastatic sites. These clones exhibit unique and stable morphological signatures, coupled with divergent growth and tumorigenic properties *in vitro*. Therefore, these findings suggest that clonal morphology as a readily assessable, practical indicator for distinguishing heterogeneous tumor populations.

### Clonal morphologic heterogeneity indicates distinct tumorigenicity and histopathology

Using the orthotopic transplantation and resection strategy, we further assessed the tumorigenic potential of these EM clones *in vivo*. Preliminary data indicated that EM2LN had relatively weak metastatic potential, EM2Lu exhibited stronger metastatic propensity, and EM6LN could form metaplastic carcinoma with lipomatous differentiation (Fig. 2A-C). Furthermore, the orthotopic tumors formed by the three EM clones exhibited the same major histological hallmarks as those of the parental EO771 orthotopic tumor (Fig. 2D-E). This indicates that the EM clones represent three distinct classes within the heterogeneity of the EO771 cell line. The tumors at the EM2LN and EM2Lu metastatic sites, reshaped by their respective metastatic microenvironments, exhibit heterogeneous pathological features (Fig. 2F-G). As these results extend beyond the immediate scope of this study, they will not be discussed in detail here.

**Fig. 2.**
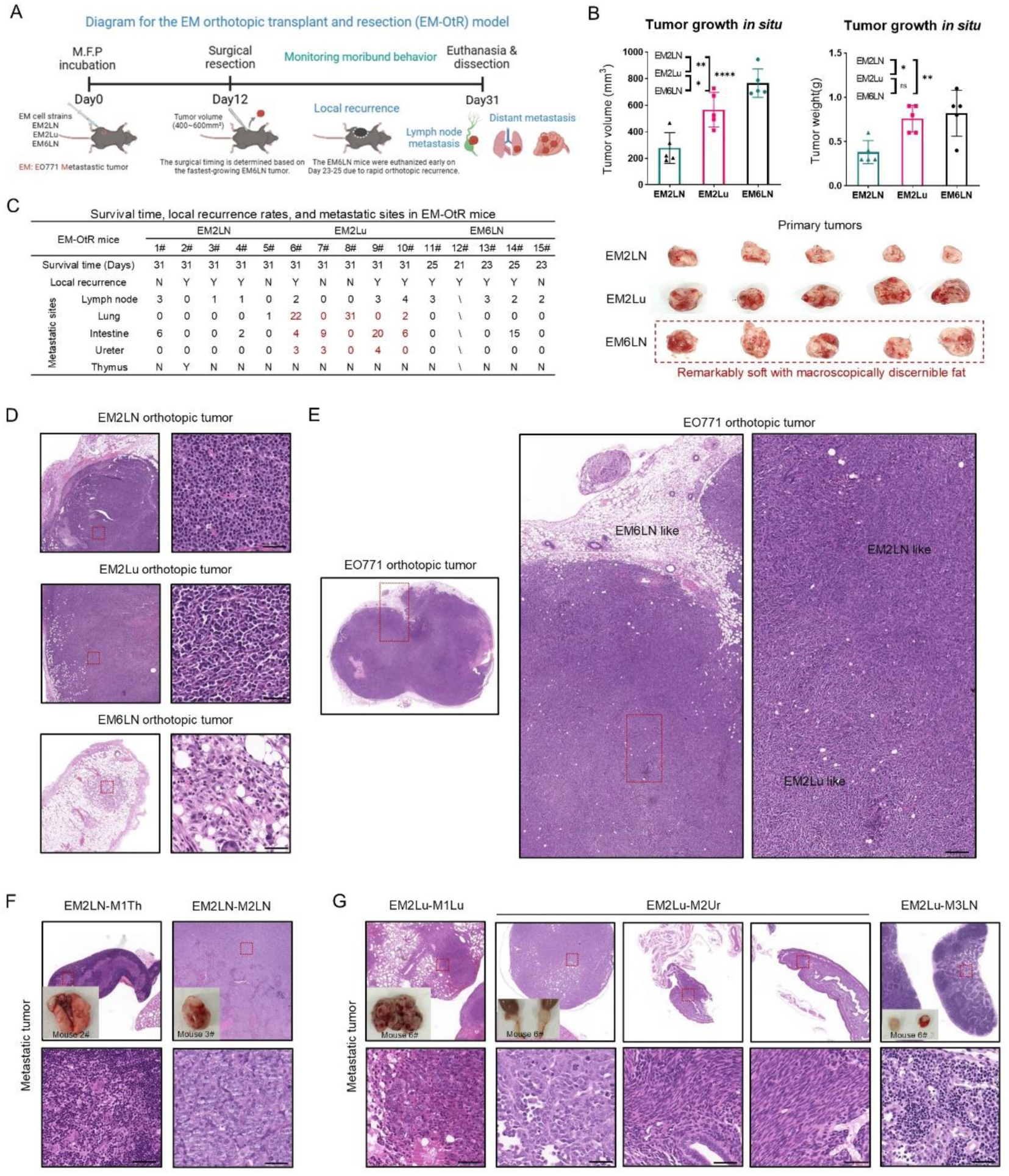
Tumorigenicity and histochemical characteristics of EM clones *in vivo*. A. Schematic diagram of the EM-OtR experimental workflow. B. Volume, weight, and representative photographs of EM orthotopic tumors. C. Statistical analysis of metastatic tumors formed by EM clones in recipient mice. D. H&E staining of EM orthotopic tumors. Scale bar: 50 μm. E. H&E staining of parental EO771 orthotopic tumors. Scale bar: 200 μm. F-G. H&E staining of tumors formed by EM2LN (F) and EM2Lu (G) at metastatic sites. “M” denotes metastatic tumor, and numbers identify individual recipient mice. Abbreviations for metastatic sites: Th, thymus; LN, lymph node; Lu, lung; Ur, ureter. Scale bar: 100 μm.

### Low-density culture induces the formation of distinct clonal morphologies in heterogeneous tumor cells

It is well established that cell lines undergo genomic alterations and transcriptomic remodeling during long-term *in vitro* culture^42,21,19,16^, suggesting that tumor cell lines themselves represent highly heterogeneous populations. To investigate whether this inherent heterogeneity can be resolved through low-density culture, we performed clonal morphology analysis on three cell lines: EO771, MDA-MB-231, and 4T1 (Fig. 3A-C). Based on morphological observations, we established several descriptive criteria: the presence of intercellular gaps within a clone, smoothness of the clone periphery, formation of spheroids indicative of escaped contact inhibition, and overall cell shape (e.g., fibroblastic, epithelial, elongated, mesenchymal). Our analysis revealed that each cell line comprised 4-5 distinct subpopulations (Fig. 3A-C), demonstrating that low-density culture can induce heterogeneous tumor cells to form unique clonal morphologies.

**Fig. 3.**
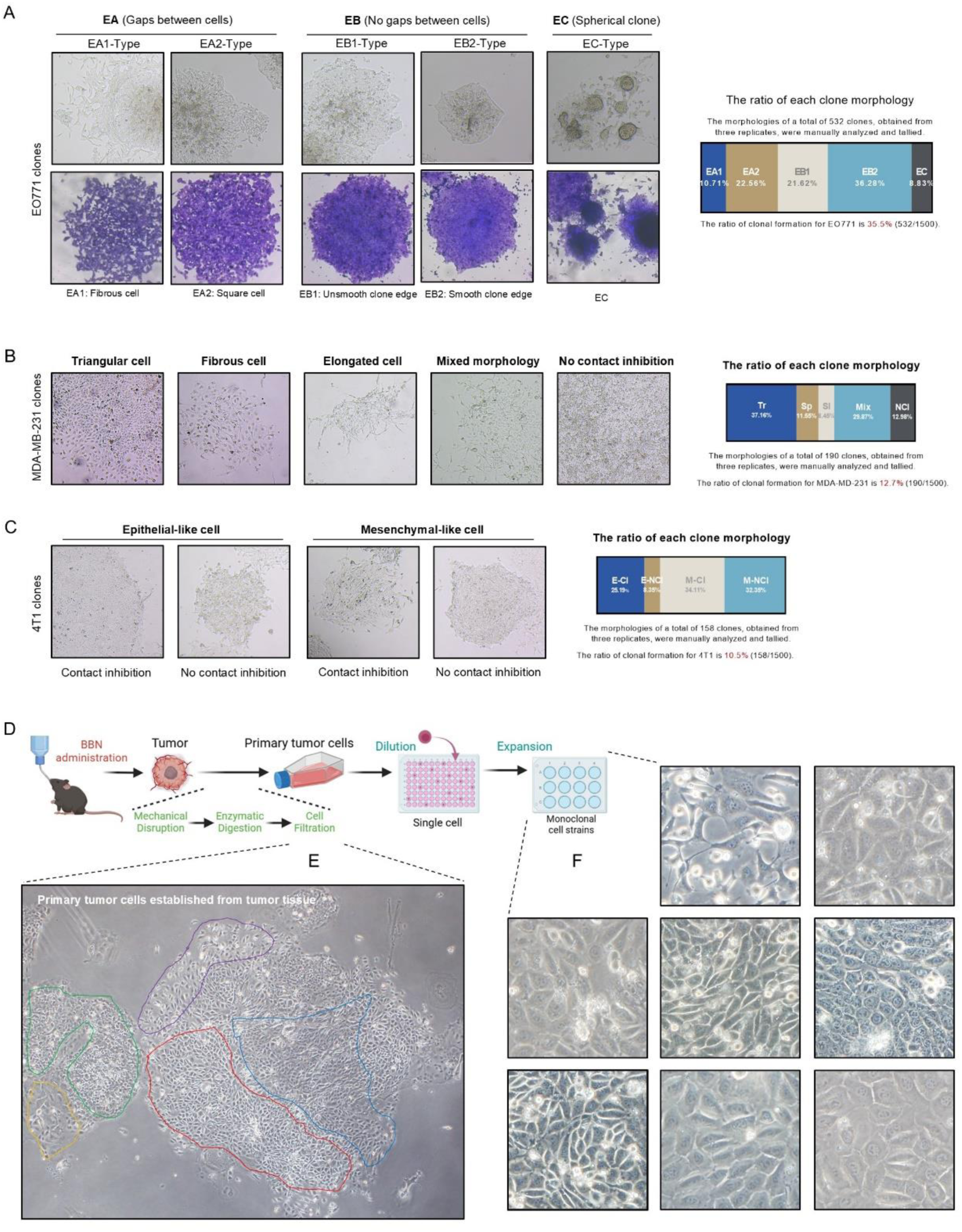
Unique clonal morphologies induced by low-density culture facilitate the identification of heterogeneous tumor cell subpopulations. A-C. Representative clonal morphologies formed by EO771 (A), MDA-MB-231 (B), and 4T1 (C) cells under low-density culture conditions. Clones derived from EO771 and MDA-MB-231 cells were classified into five distinct morphological categories, whereas 4T1-derived clones were categorized into four. Scale bar, 200μm. D. Schematic illustration of the BBN-induced bladder cancer model and the subsequent establishment of primary tumor cell cultures. E. Representative images demonstrating the heterogeneous clonal morphologies of primary tumor cells. Cell populations with distinct morphological features are outlined in different colors. Scale bar, 200μm. F. Representative images showing the morphological characteristics of 8 single-cell-derived clonal cell lines. Scale bar, 20μm.

During tumorigenesis, the accumulation of genetic variations generates highly heterogeneous cell populations. To analyze cellular heterogeneity in primary tumors, we established a chemically induced bladder cancer model and cultured the primary tumor cells *in vitro* (Fig. 3D). During culture, these primary cells exhibited diverse clonal morphologies (Fig. 3E). Furthermore, single-cell-derived sublines established from this population also displayed unique morphological features (Fig. 3F). These results confirm that low-density culture can unmask the heterogeneous composition of primary tumors through distinct clonal morphologies. Although the diversity of morphologies captured by low-density culture is more limited compared to the exhaustive profiling achieved by single-cell subcloning, it offers significant advantages in simplicity and speed for heterogeneity assessment. This practical efficiency may render the low-density clonal morphology assay particularly suitable for clinical translation in contexts such as drug sensitivity testing.

### Unique clonal morphologies manifest distinct drug response profiles

Tumor heterogeneity is a major contributor to therapeutic failure and recurrence. Theoretically, identifying clone-specific drug sensitivity profiles and vulnerabilities would enable the rational design of drug combinations targeting distinct tumor clones, thereby preventing relapse and metastasis driven by specific subpopulations.

Current drug sensitivity databases predominantly rely on high-dose, short-term (1-3 day) treatments, using cell viability to determine IC50 values. While advantageous for high-throughput screening, this strategy has limitations. Short-term, high-concentration exposure often measures acute cellular stress responses rather than mechanism-based, sustained anti-tumor effects. For instance, many chemotherapeutics exert cytotoxic effects by arresting the cell cycle—a process that can exceed 2-3 days for full effect. Consistent with this, our comparison of 2-day versus 5-day treatments of EO771 cells with gemcitabine (GEM) or MK-2206 showed that the 5-day regimen achieved equivalent inhibition at 2- to 10-fold lower doses (Fig. 4A-B).

**Fig. 4.**
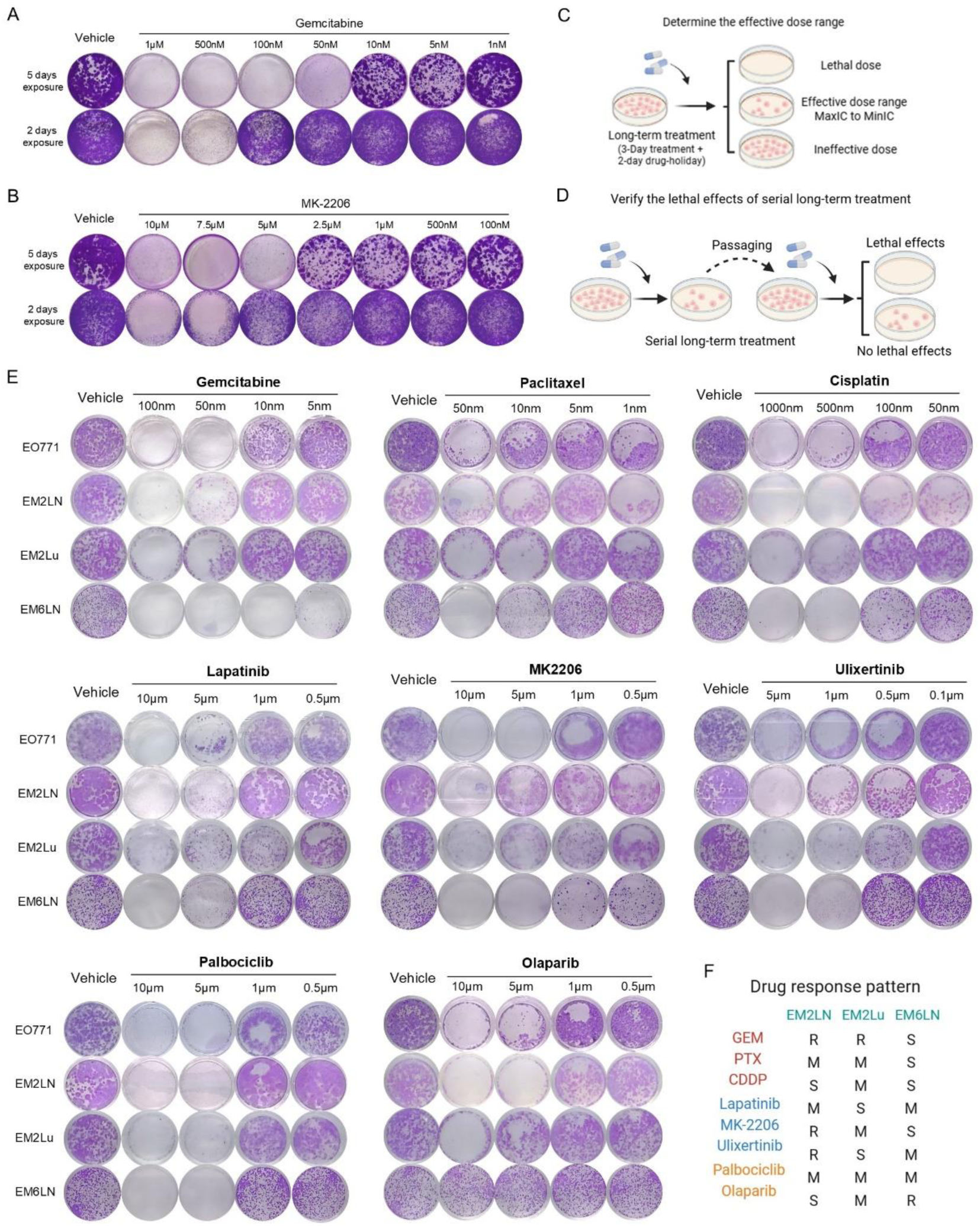
Heterogeneous tumor cell clones possess distinct drug response profiles. A-B. Proliferation inhibitory effects of long-term (5-day) versus short-term (2-day) treatment with the chemotherapy drug gemcitabine (A) and the targeted drug MK-2206 (B). The parental EO771 cell line was used to determine the effective inhibitory dose range for each drug. C. Schematic illustrating the analysis of the effective inhibitory dose range following long-term drug treatment. This range is defined as spanning from the maximum inhibitory concentration (MaxIC) to the minimum inhibitory concentration (MinIC). D. Schematic outlining the assessment of drug-induced lethal effects. Cells treated within the effective inhibitory dose range were evaluated for their ability to proliferate upon re-plating; a lack of subsequent growth indicates a cytotoxic (lethal) effect rather than a cytostatic (growth-inhibitory) effect. E. Inhibitory effects of eight drugs, each applied within its effective dose range, on the parental EO771 line and the three EM clones. F. Drug response profiles of the three EM clones relative to the parental EO771 line across the eight-drug panel.

Therefore, we optimized our drug sensitivity assay with three key modifications: First, we extended the treatment duration to 5 days (instead of 2) and assessed anti-proliferative effect rather than acute cytotoxicity. Second, we evaluated efficacy using the effective inhibitory dose range rather than a single IC50 value (Fig. 4C). Third, we distinguished between inhibitory effects and lethal effects. Drugs causing irreversible cell death upon continuous treatment within their effective dose range were defined as having a lethal effect and were excluded (Fig. 4D).

To profile anti-breast cancer drug responses, we assembled an initial panel of 22 clinical or investigational agents as breast cancer drug panel, including chemotherapy drugs, kinase inhibitors (signaling pathway), and other targeted drugs. The parental EO771 cell line was first profiled to exclude agents to which it was inherently tolerant or which exhibited lethal effects upon continuous treatment, such as Vincristine (VCR) and Dinaciclib (Fig. S1). This filtering yielded a final set of eight agents for profiling the EM clones (Fig. 4E): the chemotherapy drugs Gemcitabine (GEM), Paclitaxel (PTX), and Cisplatin (CDDP); the kinase inhibitors Lapatinib, MK-2206, and Ulixertinib; and other targeted drugs Palbociclib and Olaparib. Consequently, profiling with this refined panel revealed clone-specific response patterns (Fig. 4F): EM2LN was sensitive to CDDP and Olaparib; EM2Lu was sensitive to Lapatinib and Ulixertinib; and EM6LN was sensitive to GEM, PTX, CDDP, and MK-2206.

The Olaparib-resistant phenotype of our metaplastic-potent EM6LN model corroborates the reported intrinsic resistance in another metaplastic carcinoma model^43^, adding weight to the reliability and generalizability of our drug sensitivity results.

### Clone-sensitive drugs exert selective inhibitory effects on tumor clones

If distinct clones can be selectively eliminated by their respective sensitive drugs, then drug combinations comprising these agents hold promise for overcoming resistance caused by heterogeneous tumor cell clones. To validate the clone-selective inhibitory effect, we performed sequential GFP competitive co-culture assays. We established EM2Lu cells stably expressing GFP, mixed them with each of the three EM clones, and monitored the GFP ratio during long-term drug treatment (Fig. 5A). The results confirmed the specific sensitivities: EM2LN to olaparib, EM2Lu to ulixertinib, and EM6LN to gemcitabine (GEM) (Fig. 5B).

**Fig. 5.**
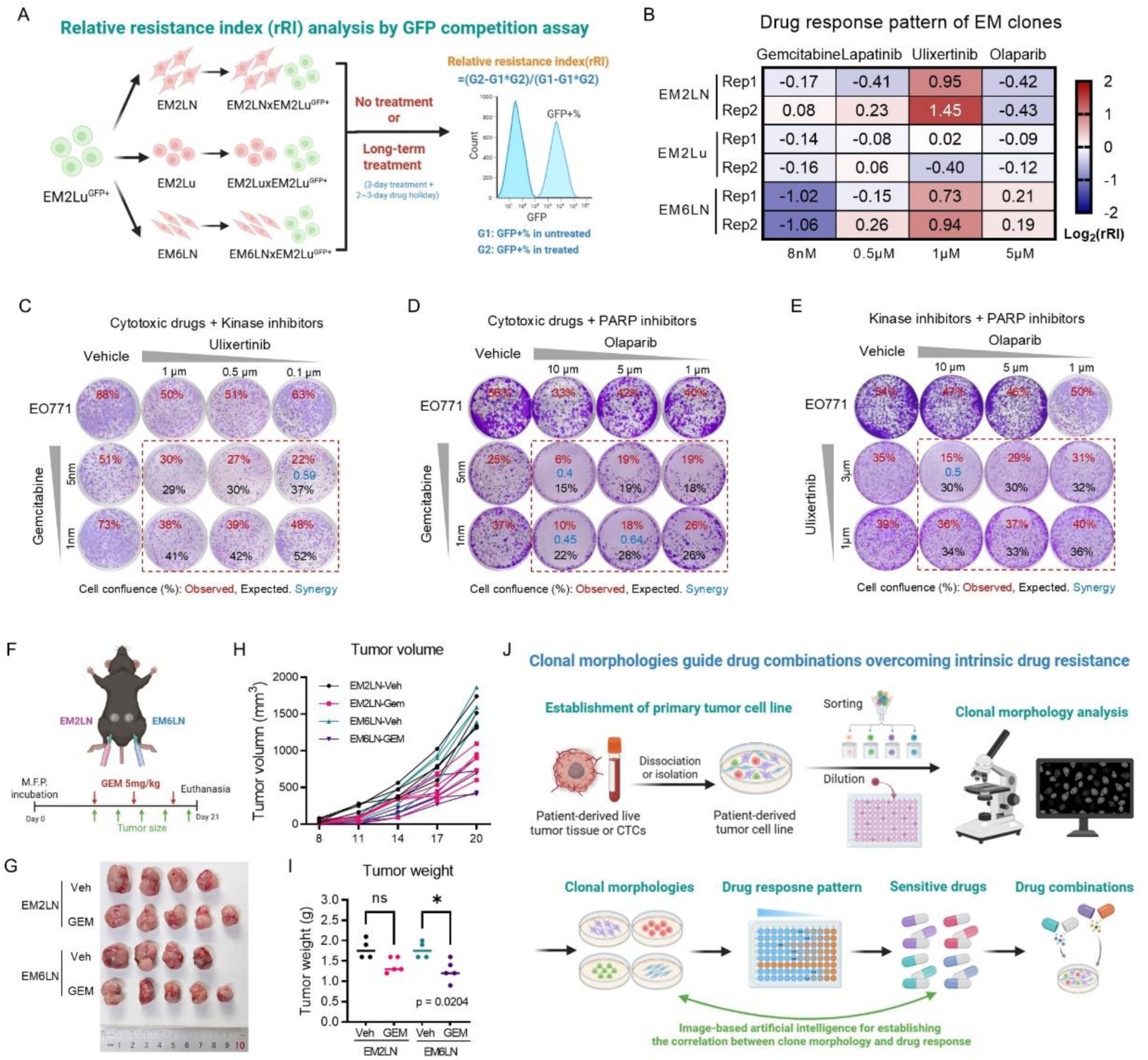
Clone-sensitive drugs selectively inhibit the proliferation of target clones. A. Schematic of the GFP competitive co-culture assay to assess the relative drug sensitivity of EM clones. EM2Lu clone was stably labeled with GFP and co-cultured with unlabeled target clones under drug treatment. B. Drug response profiles of the EM clones obtained from two independent drug sensitivity analyses, confirming clone-specific vulnerabilities. C-E. Evaluation of pairwise drug synergy for the three clone-sensitive agents. Relative cell confluence (drug-treated vs. vehicle control) was used to measure proliferation inhibition. "Observed" values represent the measured relative confluence for the combination, while "Expected" values are calculated as the product of the relative confluence values of individual drugs. A synergistic effect is indicated when Observed < Expected, with the degree of synergy quantified as Observed/Expected. F. Schematic of the *in vivo* experiment analyzing the differential response of EM2LN and EM6LN tumors to gemcitabine (GEM) treatment. G. Tumor growth curves for EM2LN and EM6LN tumors in vehicle- and GEM-treated groups. H. Images of excised tumors at the experimental endpoint. I. Tumor weight for EM2LN and EM6LN tumors in vehicle and GEM treatment groups. J. The workflow of a personalized tumor drug sensitivity assessment strategy guided by clonal morphology for drug combination discovery.

We then assessed potential synergistic effects among these three drugs in the parental EO771 cells. Most pairwise combinations showed negligible synergy (Fig. 5C-E), with only Olaparib and GEM demonstrating synergistic effects across multiple dose combinations (Fig. 5D).

Furthermore, to evaluate the clone-selective inhibitory effect *in vivo*, we orthotopically inoculated EM2LN and EM6LN clones into contralateral fat pads of recipient mice and treated them with GEM (Fig. 5F). The results showed that tumors formed by EM6LN exhibited a greater reduction in both volume and weight compared to EM2LN tumors after GEM treatment (Fig. 5G-I), indicating higher sensitivity of EM6LN to GEM.

Collectively, these results indicate that clone-sensitive drugs act by selectively inhibiting their specific subpopulations. Thus, rationally designed drug combinations informed by clonal heterogeneity can overcome resistance primarily by co-targeting and eliminating distinct subpopulations, rather than relying on synergistic drug interactions. Hence, we propose a new paradigm centered on clonal morphology to guide the identification of clone-specific vulnerabilities and drug combinations for personalized tumor therapy (Fig. 5J). The key steps are: (1) Establishing a patient-derived tumor cell culture from a surgical specimen or peripheral blood sample. (2) Generating monoclonal cell lines via single-cell sorting or limiting dilution. (3) Analyzing the clonal morphological features of these lines and grouping the heterogeneous monoclonal populations accordingly. (4) Performing drug sensitivity profiling on 1-2 representative cell lines from each morphological group. (5) Clone-specific sensitive drugs are combined to create a personalized drug cocktail designed to overcome clonal heterogeneity-driven intrinsic resistance.

### Transcriptional profiles fail to explain drug response but hint at histogenesis in metaplastic carcinoma

A pervious study has identified non-genetic factors, such as transcriptional profiles, rather than genetic mutations, as key drivers of tumor cell heterogeneity^22^. In an effort to elucidate the specific mechanisms underlying the differential drug responses among the three EM clones, we analyzed the transcriptional profiles of the EM clones compared to the parental EO771 line. Compared to the parental EO771 cells, all three EM clones exhibited substantial transcriptomic alterations, with a high number of differentially expressed genes (DEGs) among themselves (Fig. 6A). To decipher the biological significance of these EM clone-specific transcriptional profiles, we categorized the DEGs into two groups: "Activated/Suppressed" or "Opened/Closed", based on their expression levels (Fig. 6B). A DEG was classified as "Activated" or "Suppressed" if its TPM was ≥ 1 in both EO771 and an EM clone, and as "Opened" or "Closed" if its TPM was ≥ 1 in only one group. Genes with TPM < 1 in both groups were excluded from subsequent analysis. Pathway enrichment analysis was then performed on these two categories. For the "Activated/Suppressed" DEGs, EM2LN and EM2Lu displayed similar enriched pathways (Fig. 6C-D), which differed considerably from those in EM6LN (Fig. 6E). Specifically, ribosome biogenesis genes were activated, while extracellular matrix genes were suppressed in EM2LN and EM2Lu. Notably, among the "Opened/Closed" DEGs, EM6LN showed a greater number of enriched pathways. These included the "negative regulation of fat cell differentiation" pathway, which was in a "Closed" state. Five genes—Dlk1, Wnt5a, Mkx, Msx2, and Zfpm2—were enriched in this pathway (Fig. 6F). These results suggest that EM6LN possesses a transcriptional foundation that favors adipocytic differentiation, providing a partial explanation for its propensity to form metaplastic carcinoma with lipomatous differentiation *in vivo*.

**Fig. 6.**
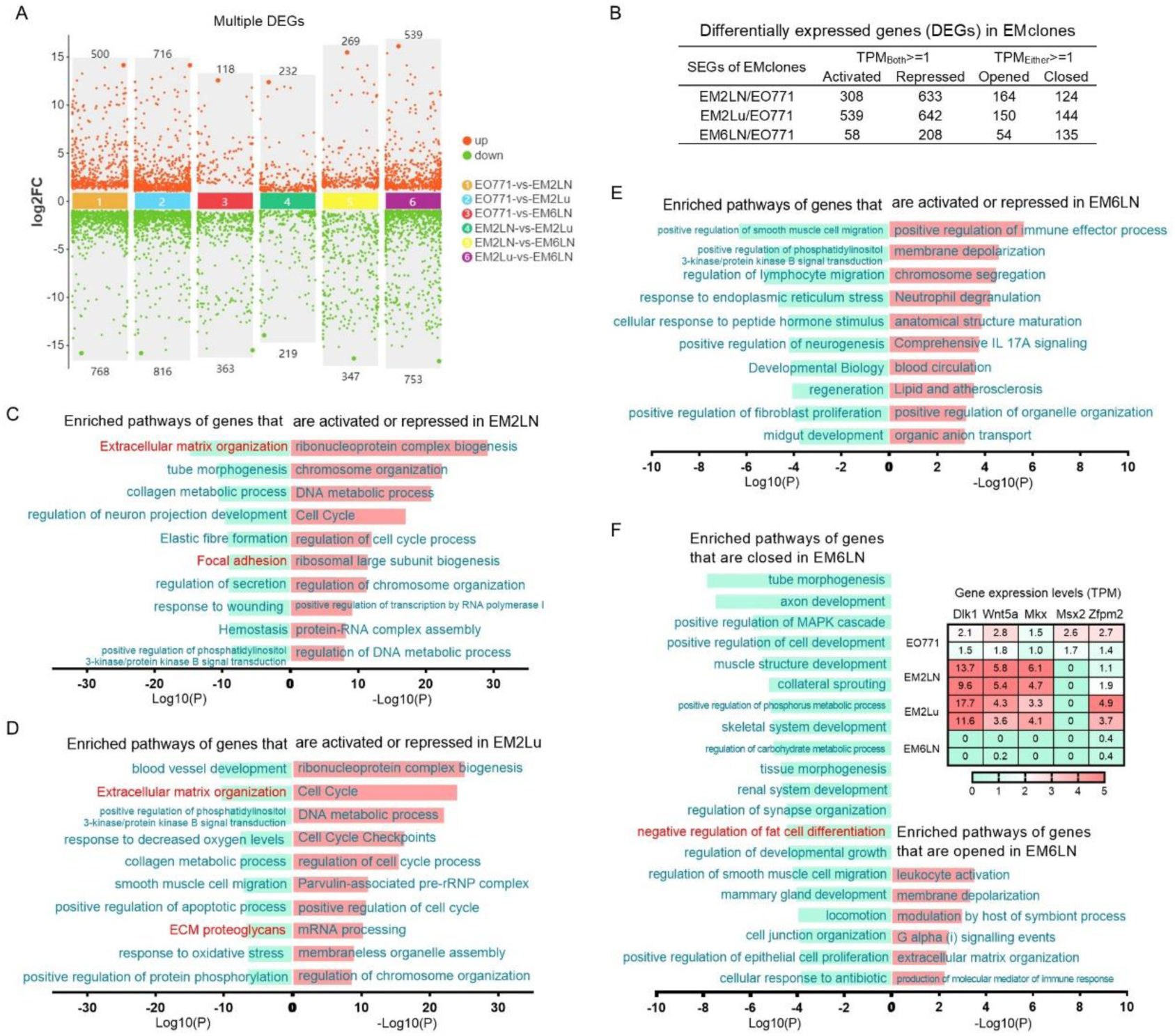
Transcriptomic profiling of EM clones. A. Number of multi-group differentially expressed genes (DEGs). DEGs were identified using an FDR threshold of < 0.05 and an absolute fold change cutoff of ≥ 2. B. Statistics of DEGs classified as “Activated/Suppressed” or “Opened/Closed”. The biological significance of a DEG was defined as “Activation” or “Suppression” if the gene’s TPM was ≥ 1 in both parental EO771 and the EM clone. It was defined as “Opening” or “Closing” if TPM ≥ 1 in only one group. Genes with TPM < 1 in both groups were excluded. C-E. Enriched signaling pathways for activated/suppressed genes in EM2LN (C), EM2Lu (D), and EM6LN (E). F. Enriched signaling pathways for opened/closed genes in EM6LN.

As a multifactorial phenotype driven by genetic and epigenetic changes^12,13,26^, drug response is difficult to correlate with transcriptional profiles based solely on differential expression. Transcriptional profiles are insufficient to explain drug-response profiles, yet they hint at the histogenesis of metaplastic carcinoma, laying important groundwork for future research into their tissue origins, which is beyond the scope of this study.

Consequently, we posit that phenotypic readouts—such as clonal morphology—may hold greater functional relevance for drug efficacy evaluation than the identification of individual molecular biomarkers (e.g., specific mutations or expression changes), as they potentially integrate the net effect of diverse underlying mechanisms.

## Discussion

Tumor cell morphological plasticity is an easily observable phenomenon. This study demonstrates that clonal morphology serves as a practical biomarker for tumor heterogeneity and reveals clone-specific drug vulnerabilities. By employing an optimized, long-term drug sensitivity assay to capture true growth inhibition, we provided proof-of-concept validation that clone-specific sensitive drugs can selectively eliminate their corresponding target clones.

A central aspect of this study is the role of culture density in revealing morphological plasticity. Tumor cells vary in their reliance on contact-mediated signaling, spanning from contact-inhibited to adhesion-dependent phenotypes. We hypothesize that high density promotes proliferative signals that mask distinct morphologies, while low density, by reducing cell contact, shifts the balance toward differentiation and unmasking of intrinsic morphological signatures. It has been reported that certain cell culture parameters, such as medium volume^44^, can reshape tumor cell behavior through specific mechanisms like hypoxia. Recent studies also have identified a minor subset of highly plastic cancer cells in models such as lung cancer, which are capable of reversibly differentiating into diverse states within the tumor^40,41^. Although we also observed differences in tumorigenicity among heterogeneous clones, our study limits itself to proposing clonal morphological plasticity as a tool for profiling drug sensitivity, as such, does not encompass an exploration of its mechanistic basis.

Current clinical approaches to tumor drug sensitivity analysis primarily rely on short-term *ex vivo* culture and testing of tumor tissue fragments or explants. Although patient-derived tumor organoids can also be used for drug testing^45,46^, the *in vitro* construction process may selectively enrich certain cell populations, thereby reducing the inherent tumor heterogeneity. Furthermore, organoid-based drug sensitivity assays are often associated with high costs and prolonged timelines, limiting their utility for large-scale *in vitro* drug screening and translational application. In contrast, our strategy requires only 1-2 weeks to establish heterogeneous primary tumor cell cultures from tissue samples and an additional 1-2 weeks to isolate distinct clonal populations. The overall cost of cell culture is low, enabling high-throughput profiling of drug response. The drug response profiling itself can be completed in approximately one week.

Several studies have demonstrated that functional precision medicine (FPM), based on direct *ex vivo* drug sensitivity testing, holds more immediate clinical translational potential compared to treatment decisions based on molecular profiling of genetic alterations^6–8^. We present a strategy that begins with discriminating heterogeneous clones by morphology, identifies their specific drug vulnerabilities, and culminates in the design of rational combinations. By placing tumor clonal heterogeneity at the center of drug sensitivity analysis, this framework establishes a new paradigm for guiding personalized precision therapy. Further development of this strategy should also encompass preclinical modeling of resistance. Selected drugs could be applied *in vitro* to generate resistant derivatives from sensitive clones; re-profiling these derivatives would thereby enable the proactive design of sequential or combinatorial regimens to counter resistance. Furthermore, while primary tumor analysis is standard, drug sensitivity testing on metastatic biopsies is strongly recommended, as the distinct microenvironment of these sites critically shapes clonal evolution and therapeutic response.

Although clonal morphology provides valuable insights, it has inherent limitations as a standalone marker of heterogeneity. Tumor cell populations exhibiting similar morphology can still be heterogeneous, harboring diverse genetic alterations and drug sensitivities. Thus, future efforts should focus on developing and integrating additional, easily measurable phenotypic biomarkers to complement morphological analysis for a more robust delineation of tumor clonal heterogeneity.

In summary, our integrated strategy presents a streamlined pipeline from morphological identification of heterogeneous clones to the design of clone-sensitive drug combinations. Its practical advantages—including low cost, rapid turnaround, reliable output, and suitability for serial assessment—collectively highlight its strong potential for clinical translation.

## Methods and materials

### Cell lines

The EO771 cell line was purchased from Xianxiang. The 4T1 cell line was obtained from the Cell Bank of Type Culture Collection of Chinese Academy of Sciences. The MDA-MB-231 cell line was maintained in-house. The EM subclones (EM2LN, EM2Lu, EM6LN) were established from EO771-derived tumors. The mouse primary bladder cell lines and their single-cell subclones were derived from BBN (N-butyl-N-(4-hydroxybutyl)nitrosoamine)-treated C57BL/6 mice. All cell lines were cultured at 37°C with 5% CO₂ in medium supplemented with 10% fetal bovine serum (FBS), 100 U/mL penicillin, and 100 μg/mL streptomycin. Specifically, 4T1 cells were grown in RPMI 1640 medium, whereas all other lines were cultured in DMEM. All cell lines and subclones were authenticated by STR profiling and/or RNA sequencing. A PCR-based assay was used to test for mycoplasma contamination in cell cultures every two weeks.

### Generation of mouse tumor models and primary tumor-derived cells

The orthotopic transplant and resection (OtR) strategy was employed to simulate post-operative recurrence and metastasis. This study used an experimental female breast cancer model, employing exclusively female mice to maintain gender-specific relevance. C57BL/6 mice were purchased from Laboratory Animal Research Center in Jiangsu University. Briefly, 1×10⁵ EO771 cells or 5×10⁵ EM clones in 50 μL DPBS were injected into the 4th mammary fat pad of anesthetized mice (2.5% avertin, 180-220 μL). Tumor volume (calculated as (L×W²)/2) was monitored until reaching 400-600 mm³, at which point the primary tumor was surgically resected. Mice were monitored daily and euthanized upon showing severe signs of disease progression (e.g., dyspnea, hunched posture) for final metastasis assessment.

A bladder cancer model was established by administering the carcinogen N-butyl-N-(4-hydroxybutyl)nitrosamine (BBN) in the drinking water. Specifically, 6- to 8-week-old C57BL/6 mice received 0.05% BBN in their drinking water, which was refreshed every five days. After approximately 5–6 months, mice presenting severe symptoms of disease progression were euthanized and tissues were collected.

Tumor tissues were aseptically dissected and minced into small fragments. To generate single-cell suspensions for initial culture (designated as passage 0, P0), the fragments were enzymatically digested according to tumor type: EO771 metastatic tumors were digested with 1.5 mg/ml collagenase type IV on a shaker for 2–3 hours, while BBN-induced bladder tumors were digested with 0.05% trypsin on a shaker for 30 minutes. After 48–72 hours of culture, the cells were digested again with trypLE. Any undigested cell clusters were removed by filtration through a cell strainer. The resulting single-cell suspension was re-plated and cultured, which was recorded as passage 1 (P1).

All mice were housed in a specific pathogen-free environment at Laboratory Animal Research Center in Jiangsu University and treated in strict accordance with protocols, which were approved by the Animal Care and Use Committee of Laboratory Animal Research Center, Jiangsu University.

### Colony formation assay and clonal morphology analysis

The colony formation assay was performed by seeding 500 cells per well in 6-well plates. During the culture period, the morphology of developing colonies was regularly monitored and documented using phase-contrast microscopy. After 8 days (or longer for extended assays), cells were fixed and stained with crystal violet for visualization and quantification.

Clonal morphology was categorized based on defined criteria that included: (1) intercellular gaps within colonies, (2) colony periphery smoothness, (3) spheroids formation (indicative of escaped contact inhibition), and (4) overall shape (e.g., fibroblastic, epithelial, elongated, mesenchymal). These features were then classified and quantified for each cell line through combined ImageJ and manual analysis.

### H&E staining and imaging

Tumor tissues were fixed in 10% formalin overnight, transferred to 70% ethanol, and paraffin-embedded following routine procedures. Subsequently, 8 μm-thick sections were prepared and stained with H&E. Whole-slide scans were acquired with a Pannoramic MIDI scanner (3DHISTECH) and analyzed using CaseViewer software (3DHISTECH).

### Chemicals

Chemotherapy drugs: 5-Fluorouracil (5-FU, S1209), Gemcitabine (GEM, S1714), Cytarabine (Ara-C, S1648), Methotrexate (MTX, S1210), Paclitaxel (PTX, S1150), Vincristine (VCR, S1241), Topotecan (TPT, S1231), Doxorubicin (DOX, S1208), Cisplatin (CDDP, S1166), Oxaliplatin (OXA, S1224), were purchased from Selleck, USA.

Kinase inhibitors and other targeted drugs: Lapatinib (S1028), Pictilisib (S1065), MK-2206 (S1078), Rapamycin (S1039), Caffeic Acid Phenethyl Ester (CAPE, S7414) Sorafenib (S1040), Ulixertinib (S7854), Trametinib (S2673), Palbociclib (S1116), Dinaciclib (S2768), Olaparib (S1060) and Selinexor (S7252) were purchased from Selleck, USA.

Most chemicals were dissolved in dimethyl sulfoxide (DMSO, V900090, Merck, USA) and to concentrations of 10mM or 100μM and aliquoted and stored at -20°C. CDDP and OXA were dissolved in Dimethylformamide (DMF, D4551, Merck, USA) and stored at -80°C. For the *in vivo* treatment experiment, gemcitabine HCl (S1149) was dissolved in saline and administered to the mice via intraperitoneal injection. Working concentrations for all chemicals were determined by inhibitory dose tests.

### Drug treatment and inhibition assay

To evaluate drug-induced growth inhibition, cells were treated for 5 days (long-term) or 2 days (short-term). For the 5-day assay, cells were seeded at 2,000 cells/well in 24-well plates; for the 2-day assay, a higher density of 15,000 cells/well was used. In both cases, treatment started 48 hours post-seeding, followed by crystal violet staining at the end of the treatment period.

The procedure for drug combination treatment was similar to that for single drugs, with one adjustment: each drug in the combination was used at twice its standard working concentration. Consequently, the mixture was applied in half the volume used for monotherapy, delivering the intended dose of each component.

### GFP competitive co-culture assay

The GFP competitive co-culture assay was performed as followings. The mixed cell populations (4,000 cells per well) were seeded into 12-well plates, with the vehicle control group receiving one-third of this cell density. Drug treatment was initiated two days post-seeding. After three days of drug exposure, the media was replaced. Tumor cells were allowed to grow for an additional two days before being harvested for passaging and flow cytometry analysis of GFP ratios. The vehicle control group was passaged at a 5:1 ratio, while the passaging ratio for drug-treated groups was adjusted based on cell counts per well. Following two days of growth post-passaging, a second round of drug treatment and GFP ratio analysis was conducted.

During cell harvesting and passaging, a small aliquot of cells was collected for GFP ratio analysis. After staining with propidium iodide (PI) for 10 minutes, flow cytometry was performed to analyze the GFP ratio in viable (PI-negative) cells. Following two rounds of drug treatment, the GFP ratios of vehicle control and drug-treated groups were used to calculate the resistance index (RI), which was employed to evaluate the impact of genetic manipulations on therapeutic response of cells to tested drugs. RI=(G1-G1*G2)/(G2-G1*G2). G1=100%-NonG1. NonG1, GFP% in vehicle control. G2=100%-NonG2. NonG2, GFP% in drug-treated.

All flow cytometry analyses were conducted using the CytoFlex instrument (Beckman) and the data were thoroughly analyzed utilizing CytExpert software (Beckman).

### Transcriptome Bulk-Sequencing

Cells were preserved in Trizol at -80°C for RNA extraction. Two biological replicates were employed for each sample. Total RNA was extracted using Trizol reagent kit (15596018, Invitrogen) according to the manufacturer’s protocol. The total RNA samples were transported on dry ice to Gene Denovo Biotechnology Company (Guangzhou, China) for library construction and bulk-sequencing (Illumina Novaseq6000). Differentially expressed genes analysis was performed by DESeq2 software between two different groups. The genes with the parameter of false discovery rate (FDR) below 0.05 and absolute fold change≥2 were considered differentially expressed genes (DEGs). Pathway enrichment analysis enrichment of DEGs were performed with Metascape (https://metascape.org).

### Statistical analysis

The statistical methods employed were tailored to the specific type of experiment, including log-rank (Mantel–Cox) tests, as detailed in the respective figure legends. Data were presented as mean ± SD. Statistical significance was determined by P values less than 0.05. Significance levels were denoted as follows: *P < 0.05, **P < 0.01, ***P < 0.001, and ****P < 0.0001. Statistical analyses were conducted using GraphPad Prism 9.0 software (GraphPad Software).

## Data availability

All raw data generated or analyzed throughout this study are accessible upon reasonable request from the corresponding author. The RNA-Seq data from this study have been deposited in the NCBI Sequence Read Archive (SRA) and are publicly accessible under the accession number PRJNA1268032. Further information and requests for resources and reagents should be directed to Xiaoxi Li.

## Conflict of interest

All authors declare no competing interests.

## Acknowledgments

There was no dedicated project support for this study. The authors utilized their personal resources and institutional facilities to carry out the research and complete the manuscript. Diagrams presented in this article were created with BioRender.com.

## Author contributions

Conceptualization, X.L.; Methodology, X.L. L.Liu.; Software, X.L.; Validation, X.L. and L.Liu.; Formal analysis, X.L; Investigation, X.L., L.Liu., Y.Y., L. Luo., Y.J., W.R., and M.Y.; Resources, X.L.; Data curation, X.L. and L.Liu.; Writing-Original Draft, X.L.; Writing—Review and Editing, X.L.; Visualization, X.L.; Supervision, X.L.; Project Administration, X.L.. All authors have read and agreed to the published version of the manuscript.

## Notes

### Competing Interest Statement

The authors have declared no competing interest.

