## Supplementary material for "Clonal morphology-guided combination therapies overcome heterogeneity-driven drug tolerance": Figure S1

**List of Supplementary Materials**

Supplementary Figure 1

Fig. S1. Analysis of effective inhibitory dose ranges.

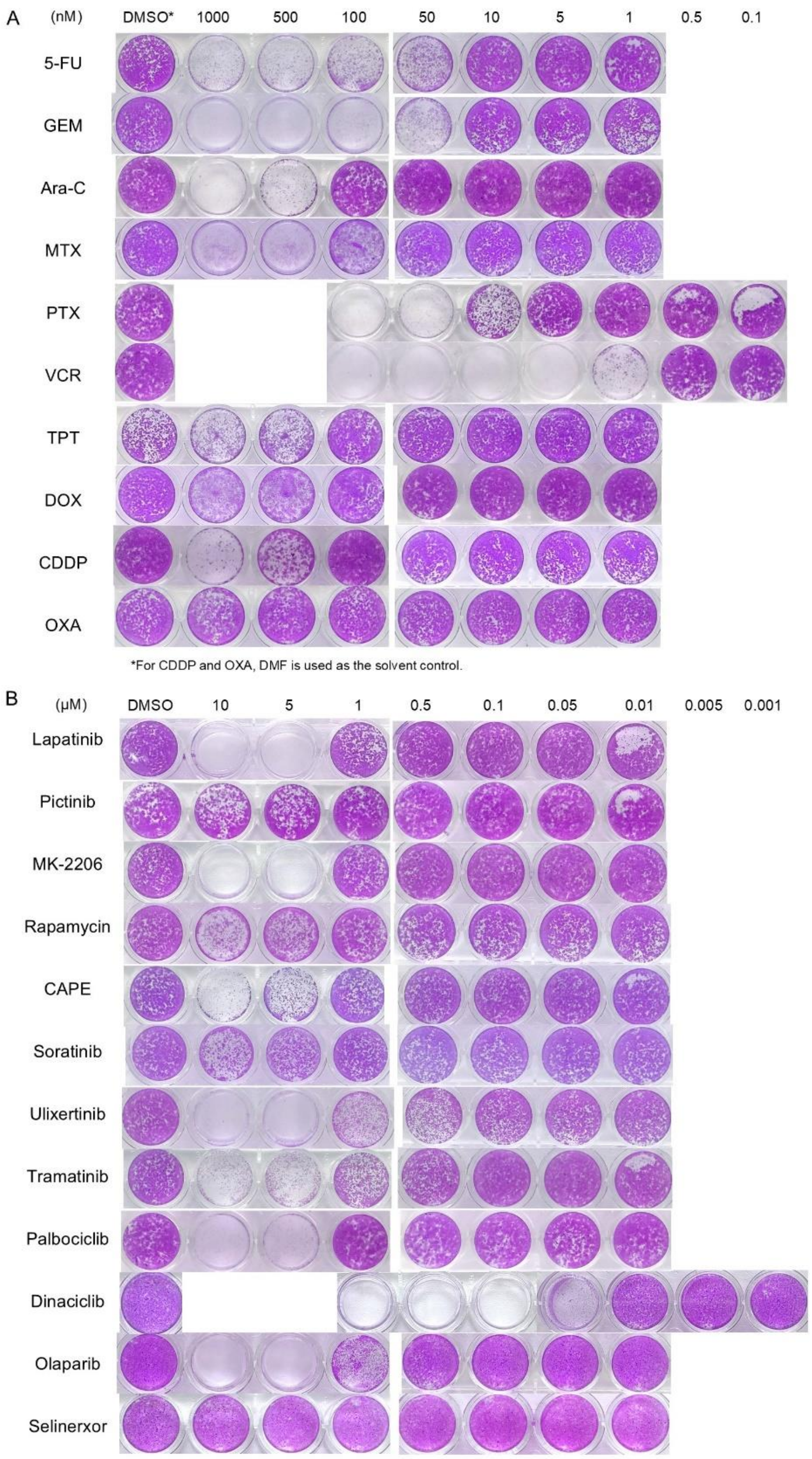


**Fig. S1. Analysis of effective inhibitory dose ranges.** A-B. Proliferation inhibition of EO771 cells following long-term treatment with a breast cancer drug panel, comprising 10 cytotoxic drugs (A) and 12 targeted agents (B).
